# An *in vivo* Probabilistic Atlas of the Human Locus Coeruleus at Ultra-high Field

**DOI:** 10.1101/2020.02.03.932087

**Authors:** Rong Ye, Catarina Rua, Claire O’Callaghan, P Simon Jones, Frank Hezemans, Sanne S. Kaalund, Kamen A. Tsvetanov, Christopher T. Rodgers, Guy Williams, Luca Passamonti, James B. Rowe

**Affiliations:** Department of Clinical Neurosciences, University of Cambridge, Cambridge, UK; Wolfson Brain Imaging Centre, University of Cambridge, Cambridge, UK; Brain and Mind Centre and Central Clinical School, Faculty of Medicine, University of Sydney, Sydney, Australia; Department of Psychiatry, University of Cambridge, Cambridge, UK; Medical Research Council Cognition and Brain Sciences Unit, Cambridge, UK; Department of Psychology, University of Cambridge, Cambridge, UK; Istituto di Bioimmagini e Fisiologia Molecolare (IBFM), Consiglio Nazionale delle Ricerche (CNR), Milano, Italy; Cambridge University Hospitals NHS Foundation Trust, Cambridge, UK

**Keywords:** locus coeruleus, probabilistic atlas, 7T MRI, neurodegenerative disorders, contrast-to-noise ratio

## Abstract

Early and profound pathological changes are evident in the locus coeruleus (LC) in dementia and Parkinson’s disease, with effects on arousal, attention, cognitive and motor control. The LC can be identified *in vivo* using non-invasive magnetic resonance imaging techniques which have potential as biomarkers for detecting and monitoring disease progression. Technical limitations of existing imaging protocols have impaired the sensitivity to regional contrast variance or the spatial variability on the rostrocaudal extent of the LC, with spatial mapping consistent with *post mortem* findings. The current study employs a sensitive magnetisation transfer sequence using ultrahigh field 7T MRI to investigate the LC structure *in vivo* at high-resolution (resolution 0.4×0.4×0.5 mm, duration seven minutes). Magnetisation transfer images from 53 healthy older volunteers (52-84 years) revealed the spatial features of the LC and were used to create a probabilistic LC atlas for older adults, appropriate for clinical research. Consistent rostrocaudal gradients of slice-wise volume, contrast and variance differences of the LC were observed, mirroring distinctive *ex vivo* spatial distributions of LC cells in its subregions. The contrast-to-noise ratios were calculated for the peak voxels, and for the averaged signals within the atlas, to accommodate the volumetric differences in estimated contrast. The probabilistic atlas is freely available, and the MRI dataset is available for researchers, for replication or to facilitate accurate LC localisation and unbiased contrast extraction in future studies.

## 1. Introduction

The locus coeruleus (LC) is the principal source of noradrenergic neuromodulatory inputs to the forebrain (Szabadi, 2013, Sara, 2009). The LC influences multiple functions including attentional control (Aston-Jones and Cohen, 2005), stress response and emotion (Itoi and Sugimoto, 2010), memory consolidation and retrieval (Sara, 2015), decision making (Yu and Dayan, 2005, Bouret and Sara, 2005), sleep-wake cycle (Berridge et al., 2012), and autonomic control (Samuels and Szabadi, 2008).

The human LC is a small, rod-shaped nucleus located at the dorsal boundary of the pons with a rostrocaudal extent of approximately 12 - 17 mm (Fernandes et al., 2012). It is adjacent to the lateral floor of the fourth ventricle and projects long collateralised ascending axons with minimal myelination - a property may render the LC vulnerable to environmental toxins (Braak et al., 2003, Mather and Harley, 2016). The LC neurons degenerate in many neurological disorders. For example, abnormal pre-tangle tau protein is detected in the LC in Alzheimer’s disease from the earliest stages (Braak et al., 2011, Grudzien et al., 2007, Kelly et al., 2017), and severe cell loss is observed in Alzheimer’s disease, Parkinson’s disease and Progressive Supranuclear Palsy (Theofilas et al., 2017, Surmeier et al., 2017, Kaalund et al., 2020). Dysregulation of the LC noradrenergic system has been associated with clinical symptoms and cognitive impairments in neurodegenerative disorders (Weinshenker, 2008, Kehagia et al., 2010, Passamonti et al., 2018a). Given the importance of the LC in brain function and pathogenesis, the detection and monitoring of LC degeneration *in vivo* with non-invasive imaging techniques would be valuable for presymptomatic diagnosis and experimental medicines studies of early-stage interventions (Betts et al., 2019).

Several studies have used T1-weighted sequences for the investigation of the LC in healthy adults (Keren et al., 2009, Betts et al., 2017, Tona et al., 2017). The location of hyperintense voxels in the dorsal pons corresponds to the approximate spatial distribution of noradrenergic LC cells and has been validated as LC by histology (Keren et al., 2015). The source of the LC contrast in T1-weighted (T1-w) images has been attributed to the presence of neuromelanin (NM) in pigmented LC neurons, which acts to reduce T1 and also causes magnetisation transfer (MT) effects (Chen et al., 2014, Trujillo et al., 2017, Priovoulos et al., 2018, Sasaki et al., 2006). In particular, the paramagnetic metal chelation and autophagic processes linked to the neuromelanin production create contrast in T1-w images (Sulzer et al., 2018). Recent evidence from preclinical models suggests that the LC contrast may also partially depend on the enriched water content of the LC cell bodies which would cause an increased T_1_ and relatively low MT ratio (MTR) of the LC structure relative to the surrounding tissues (Watanabe et al., 2019a, Watanabe et al., 2019b). Although the specific contrast source differs in these biophysical models, both imply an association between the LC contrast level and the fraction of LC neurons, i.e. the diminished LC contrast should reflect either reduced neuromelanin concentration or increased proton density accompanied by neuronal loss or neuroinflammation (Zucca et al., 2014, Hansen et al., 2016, Lambert et al., 2013).

The contrast to noise ratio (CNR) estimated from *in vivo* imaging has been widely used as a marker of LC structural integrity for disease stratification, differential diagnosis (Ohtsuka et al., 2013, Takahashi et al., 2015, Ohtsuka et al., 2014, Matsuura et al., 2013), or to explain variance in cognitive performance (Clewett et al., 2018, Dahl et al., 2019, Hammerer et al., 2018). Previous 3T imaging studies typically employed 2D sequences with large slice thickness to improve the signal-to-noise ratio (SNR) and reduce the scan duration. However, this approach increases partial volume effects that hinder the accuracy of imaging localisation and quantification, especially for an elongated structure like the LC where the signal is distributed along the longitudinal axis of 2D sequence voxels. An alternative MT-weighted turbo flash (MT-TFL) 3-D sequence has been developed to increase resolution and speed of LC imaging, coupled with enhanced SNR using ultra-high field 7T MRI (Priovoulos et al., 2018). This study also demonstrated that 7T MT-TFL sequence allows safe and more tolerable scans which are suitable for patients and have superior SNR and CNR relative to earlier 3T T1-w sequences.

The application of LC CNR as an imaging biomarker in clinical studies relies on an adequate and unbiased estimation of the signal in disease groups where the neuronal loss relates to volumetric reduction (Theofilas et al., 2017). Manual segmentation is often used for extracting the LC signal, but this method relies on subjective estimations that only have fair to good intra-rater reliability (Tona et al., 2017, Hammerer et al., 2018). The manual segmentation approach becomes more unreliable when the signal-to-noise is reduced by pathological changes and when the signal is more dispersed by anisotropic resolution. Some studies implemented a semi-automated, threshold-based segmentation method (Chen et al., 2014, Dahl et al., 2019). A stringent threshold was calculated from a pontine reference region, which was used to automatically extract the LC voxels within the possible location. While this semi-automatic approach provides excellent reproducibility (Langley et al., 2017), the segmentation-based CNR calculation in patient groups generally suffers from signal inflation. Based on the current hypothesis that neuromelanin deposits are the source of contrast in MT sequences, the reduction in the contrast-to-noise ratio observed in the LC in disease is likely to reflect the neuronal loss that accompanies the volumetric decrease of the LC. On this basis, the segmentation-based algorithm for LC contrast estimation only identifies signals from the surviving LC neurons where the volumetric loss is not accounted for, leading to a potentially inaccurate (and biased) estimation of LC integrity in patients.

Here we propose an alternative, atlas-based, method for extracting the LC CNR which has three main advantages studies. First, a human atlas is useful for directing the anatomical localisation and spatial definition of the LC. The cell loss confound and resulting volumetric reduction in neurodegeneration is controlled for when comparing the group differences of the CNR within the same spatially confined volume determined by the atlas. Second, the method relies on an accurate co-registration between the raw images and the atlas, which can be achieved with high-performance normalisation algorithms for cross-modality brain image registration (Klein et al., 2009). LC signals can be transformed to the standard space for consistent between-subject comparisons while the relative locations are preserved. Third, the standardised atlas enables comparable analyses and more interpretable results across studies and disease groups. We develop the atlas in the context of mid and later life, in part because of the greater neuromelanin based contrast with age, but also in anticipation of the use of the atlas to study neurodegenerative disease, where an atlas based on young adult brains would not be appropriate.

The current study aimed to: (1) produce an accurate *in vivo* mapping of the LC at high resolution using a MT-TFL sequence at ultrahigh field strength; (2) create an atlas of the human LC using a cohort that enables translational studies in neurodegenerative diseases; (3) release this 7T-based human LC atlas and the underlying dataset, to enable replication studies and/or to develop alternate strategies for future research.

## 2. Materials and Methods

To achieve a useful atlas, our criteria were threefold: (i) to include a sequence sensitive to LC signal and a normal population over 50 years, that is relevant to deficits of the LC associated with dementia and neurodegenerative disorders, (ii) to incorporate robust and accurate normalisation to a standard space to facilitate comparisons across individuals and across studies, and (iii) to produce a probabilistic atlas of the LC from which, taking in consideration the trade-off between sensitivity and specificity, researchers will be able to adaptively select their LC region of interest (ROI).

### 2.1 Participants

Fifty-six healthy adults (24 females, age range: 52 – 84 years, mean age ± SD: 66 ± 7 years) were recruited after written informed consent in two concurrent research protocols. The protocols were approved by Cambridge Research Ethics Committee (16/EE/0084 and 10/H0308/34). The exclusion criteria were: i) lack of mental capacity to consent, ii) any contraindication to ultra-high field 7T MRI, iii) any significant neurological (e.g. stroke, epilepsy, traumatic brain injury) or psychiatric (e.g. schizophrenia, bipolar disorder, current depression) or medical disorder (e.g., ischemic heart disease, cardiac rhythm abnormalities, significant non-ischemic cardiac disease, uncontrolled hypertension and known hepatic or renal failure).

Two subjects were excluded due to abnormal incidental findings on their T1-w structural images. One participant was excluded because of excessive movement during the scan. This yielded 53 completed data to be used for further analysis.

### 2.2 MRI Acquisition

The MR images were acquired with a 7T Magnetom Terra scanner (Siemens, Erlanghen, Germany). For radiofrequency transmission, a 32-channel receive and circularly polarised single-channel transmit head coil (Nova Medical, Wilmington, USA) was used. Earplugs and cushions were used to reduce the acoustic noise and movement, respectively.

We used a 3-D high-resolution MT-TFL sequence for imaging the LC (Priovoulos et al., 2018). 112 axial slices were used to cover both the midbrain and the pontine regions. The sequence applied a train of 20 Gaussian-shape RF-pulses at 6.72 ppm off-resonance, 420° flip-angle, followed by a turbo-flash readout (TE = 4.08 ms, TR = 1251 ms, flip-angle = 8°, voxel size = 0.4 x 0.4 x 0.5 mm^3^, 6/8 phase and slice partial Fourier, bandwidth = 140 Hz/px, no acceleration, 14.3%-oversampling, TA ~ 7 min). For each subject, the transmit voltage was adjusted based on the average flip angle in the central area of the pons obtained from a B1 pre-calibration scan. The MT-TFL sequence was repeated three times and next it was averaged off-line to improve the SNR. An additional MT scan was acquired with the same parameters as above but without the off-resonance pulses to directly allow the examination of the MT effect.

A high resolution isotropic T1-w MP2RAGE image was also acquired sagittally for anatomical coregistration using the UK7T Network harmonised protocol (Clarke et al., 2019): TR = 3500 ms, TE = 2.58 ms, BW = 300 Hz/px, voxel size = 0.7 x 0.7 x 0.7 mm, FoV = 224 x 224 x 157 mm, acceleration factor (A>>P) = 3, flip angles = 5/2° and inversion times (TI) = 725/2150 ms for the first/second images. The total acquisition time of the MP2RAGE was 7 minutes and 51 seconds.

### 2.3 Image Processing and Coregistration Pipeline

The Advanced Normalization Tools (ANTs v2.2.0) software and in-house Matlab (R2018b, The MathWorks, Massachusetts, United States) scripts (available upon request) were used for image pre-processing and the standardisation of MT images. Both types of MT images, with off-resonance pulses switched on (MT-on) and off (MT-off), were first corrected for spatial inhomogeneity using the N4 bias field correction function in ANTs (number of iterations at each resolution level: 50×50×30×20, convergence threshold: 1×10^−6^, isotropic sizing for b-spline fitting: 200) (Tustison et al., 2010). To further suppress the noise and compensate for potential small motion artefacts, three repeats of MT-on images were simultaneously coregistered and averaged using the antsMultivariateTemplateConstruction2 function with rigid transformation for intra-subject, intra-modality image average. The T1-w MP2RAGE data were generated offline from the complex GRE images as in (Clarke et al., 2019). T1-w skull-stripped images were obtained after tissue type segmentation and reconstruction using SPM12 (v7219). In addition to the inclusion of grey-and white-matter masks for brain extraction, the cerebrospinal fluid was also added to the brain mask to avoid excessive brain stripping.

These pre-processed MT-weighted and T1-w images were then entered into a T1-driven, cross-modality coregistration pipeline to warp the individual MT-on and MT-off images to the isotropic 0.5 mm ICBM152 (International Consortium for Brain Mapping) T1-w asymmetric template (Fonov et al., 2011). As shown in Figure 1, the individual T1-w images were first coregistered to the MT-on image with rigid only transformation. The MT-off image was used as the intermediate step for bridging the two modalities because the MT-off images shared similar tissue-specific contrasts with both T1-w MP2RAGE and MT-on images. In parallel, a study-wise T1-w structural template was created using individual skull-stripped T1-w images. Native T1-w images were rigid and affine transformed followed by a hierarchical nonlinear diffeomorphic step at 5 levels of resolutions, repeated by 6 runs to improve convergence. Max iterations for each resolution from the coarsest level to the full resolution were 100×100×70×50×20 (shrink factors: 10×6×4×2×1, smoothing factors: 5×3×2×1×0 voxels, gradient step size: 0.1 mm). The Greedy Symmetric Normalisation (SyN) was adopted for the transformation model of the deformation step (Avants et al., 2008).

**Figure 1.**
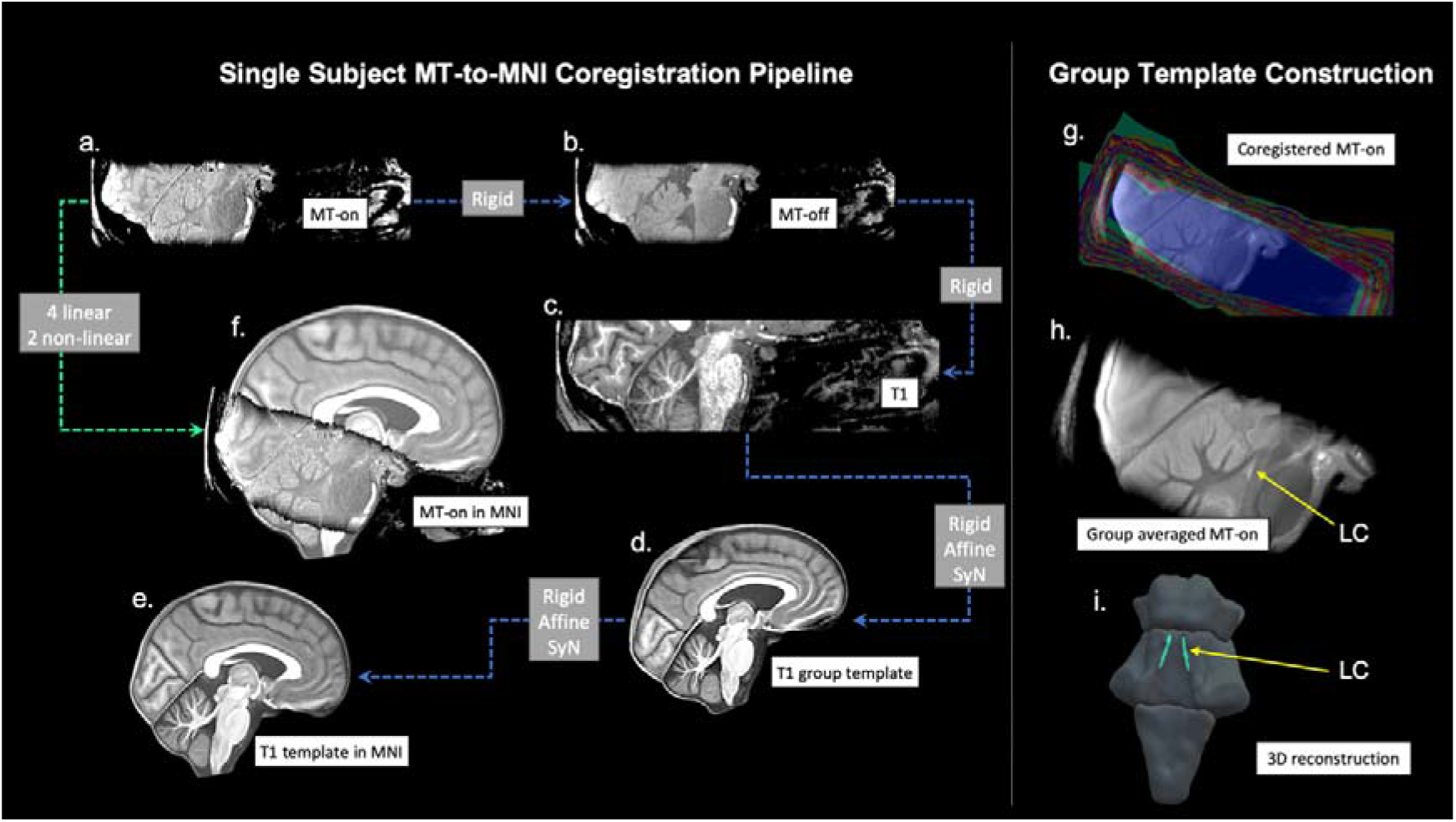
The T1-driven image registration pipeline for the normalisation and group template construction of MT images. Individual MT-on image (a) was coregistered with the MT-off image (b) and the T1-MP2RAGE image (c) using rigid-body transformation for intra-subject image registration. T1-w images from all subjects were firstly skull-stripped then used to create a study-wise template (d) which was subsequently registered with the ICBM152 standard brain (e). Linear and nonlinear transformation parameters estimated in the previous coregistrations were applied in one step to normalise individual MT-on images to the MNI space (f). Coregistered MT-on images (g) were averaged to obtain a group template for visualisation and segmentation. The location of the LC was shown on the sagittal view of the MT group template (h) and in the 3D reconstruction of the brainstem (i).

The resulting T1-w group template was then registered to the standard ICBM152 T1-w brain following the similar rigid-affine-SyN steps at 4 resolution levels (max iterations: 100×70×50×50, convergence threshold: 1×10^−6^, shrink factors: 8×4×2×1, smoothing factors: 3×2×1×0 voxels). For all the above registration steps, cross-correlation was used for similarity metrics estimation as it performs better for linear and non-linear components during intra-modality registration. Four linear transformation matrices (rigid + affine) plus two non-linear deformations were estimated from four registration steps as follows (in order): MT-off to MT-on, T1-w to MT-off, T1-w to T1-w group template and T1-w group template to ICBM152 T1-w template. These parameters were then used as the roadmap for MT images standardisation to the ICBM brain in one step. A trilinear interpolation method was selected to preserve the absolute location and relative contrast of the signal. The coregistered n=53 MT-on and MT-off images were averaged, respectively, to form the group-level MT-on and MT-off templates which will be used to facilitate visualisation and region of interest (ROI) definitions for LC segmentation in the following step. To calculate the MTR map voxel-by-voxel and compare across participants, a single MT-on image acquired immediately before the MT-off image was also normalised to the ICBM152 brain for each subject.

### 2.4 Registration Accuracy Evaluation

To evaluate the accuracy for the T1-driven registration steps from native space to standard brain space, an anatomical landmark specific to the LC and available in the automatic segmentation pipeline was extracted on native T1-w MP2RAGE images. FreeSurfer (v6.0.0) recon-all function was used for segmenting individual pons and a Bayesian segmentation algorithm (Iglesias et al., 2015) based on a probabilistic atlas of the brainstem was implemented to improve the spatial resolution (0.33 mm isotropic) and segmentation accuracy. The *highres* option was selected for submillimetre resolution structural images in the current dataset (Zaretskaya et al., 2018). The boundaries of the pons were derived from the automated segmentation then registered to the ICBM152 template by applying the transformation matrices estimated in the previous steps. Pairwise Dice Similarity Coefficients (DSC) between individual (I) and standard (S) pons were calculated with *DSC* = 2 (*I* ∩ *S*)/(*I* + *S*). A DSC closer to 1 indicates a higher level of similarity and registration accuracy, suggesting that the LC variability observed in later stages can be more confidently explained by intrinsic structural variations rather than by registration differences.

### 2.5 Reliability: Semi-automated LC Segmentation

A semi-automated threshold-based method was employed to provide more reliable outcomes than manual segmentation of the LC (Chen et al., 2014, Langley et al., 2017). In implementing our method we pre-defined three elements: a) the a *priori* rostrocaudal boundaries for the extent of the LC spatial distribution; b) the selection of a reference region used to estimate the noise and calculate the threshold; c) and the “searching area” for possible LC locations to which a threshold is applied, and from which LC voxels are extracted. The ITK-snap software was used to visualise the images and draw ROIs. The methods are described in detail below:

1. Given that boundaries of gross brainstem structures are easily identifiable on the MT-off image, the MT-off study template in MNI space was used to define the rostral and caudal anatomical limits of the LC. In keeping with previous studies that determined the spatial extent of the LC (Keren et al., 2009, Betts et al., 2017, Priovoulos et al., 2018), the rostral and caudal boundaries were respectively defined at the level of the decussation of the superior cerebellar peduncles (SCP), beneath the level of the inferior colliculi (Z = −16 mm, slice number = 112), and the recess of the 4^th^ ventricle (Z = −29 mm, slice number = 86) (Figure 2B).
2. Candidate reference regions previously used in literature were defined on the axial slices of the MNI MT-on study template because of the visibility of the white-matter structures, including the central pontine area (PT), dorsal PT, left PT, right PT, left SCP and right SCP (Figure 2C). A region was considered suitable as reference area if it showed low internal noise (high SNR) and distinctive MT effects relative to the LC (high MTR resulting in significant signal suppression in MT-on images). Considering the potential inter-hemispheric difference of PT signals, some studies have selected ipsilateral PT areas as reference regions for left and right LC segmentation (Garcia-Lorenzo et al., 2013, Betts et al., 2017, Tona et al., 2017). If the signal asymmetry were evident in all PT structures including the LC, then the ipsilateral referencing approach would have been adopted in this study. Conversely, if signals were only asymmetric for PT structures but not the LC, then the use of bilateral reference regions would have not been considered appropriate as it would have introduced a biased baseline for LC contrast estimation.
3. The searching area for the segmentation of the LC was respectively defined for the left and right LC on the study-wise MNI MT-on template where the group-level representation of the LC can be easily visualised with high-signal contrast (Figure 2D). MT-off template was overlaid onto the group averaged MT image and colour-coded with jet map to enhance the contrast between the brainstem and the 4^th^ ventricle which is less visible on MT-on images. This procedure was used to direct the positioning of the LC searching area and to avoid the contamination of signal from the voxels belonging to the 4^th^ ventricle.
4. To control for the effect of dimensionality in signal extraction, the size of the reference regions was designed to match the searching area of the LC. A specific circular ROI with 2.5 mm diameter was selected for the left and right LC, spanning the full rostro-caudal extent of the LC (13.5 mm). A ROI with the same shape and size of the LC searching area was centred at the root of the left and right SCP. For all other PT candidate reference ROIs, a 4 x 4 x 4.5 mm^3^ cubic ROI was placed at the level of the middle pons. The total volumes were matched between the circular (70.88 mm^3^) and cubic (72 mm^3^) ROIs.
5. Within the searching area, the LC voxels were segmented using a stringent threshold: *T*= *Mean*_*REF*_ + 5 × *SD*_*REF*_. Due to the enhanced SNR at 7T, a more stringent threshold relative to previous studies with similar segmentation pipeline was selected (Chen et al., 2014, Dahl et al., 2019). A binary decision was made for any given voxel in the LC area based on the threshold (T) and voxels were categorised as belonging to the LC if their intensities were greater than the mean intensity of the reference region (Mean_REF_) by more than 5 standard deviations (SD_REF_) (*vice versa* for non-LC voxels).

### 2.6 Signal Evaluation

The SNR in the LC searching area and PT candidate reference ROIs were examined on the averaged MT-on image 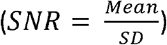. The single MT-on and MT-off images were aligned and warped to the MNI standard space. Individual MTR maps were created by voxel-by-voxel computation of the MTR as follows: 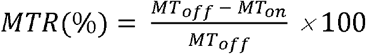. Subject-specific MTR values were next extracted for the LC searching area and PT candidate reference ROIs using individual MTR maps. In addition, mean signal intensities in the left and right LC, SCP and PT areas were extracted from the MT-on images. Paired t-tests were performed to examine the signal asymmetry between the left and right side. The ipsilateral referencing approach was only selected if lateralised signal distribution was also found in the LC. The decision regarding the appropriate reference region to use was made on the basis of the characteristic high SNR and high MTR that were sufficient for generating robust automated segmentation of the LC and reliable CNR estimation.

**Figure 2.**
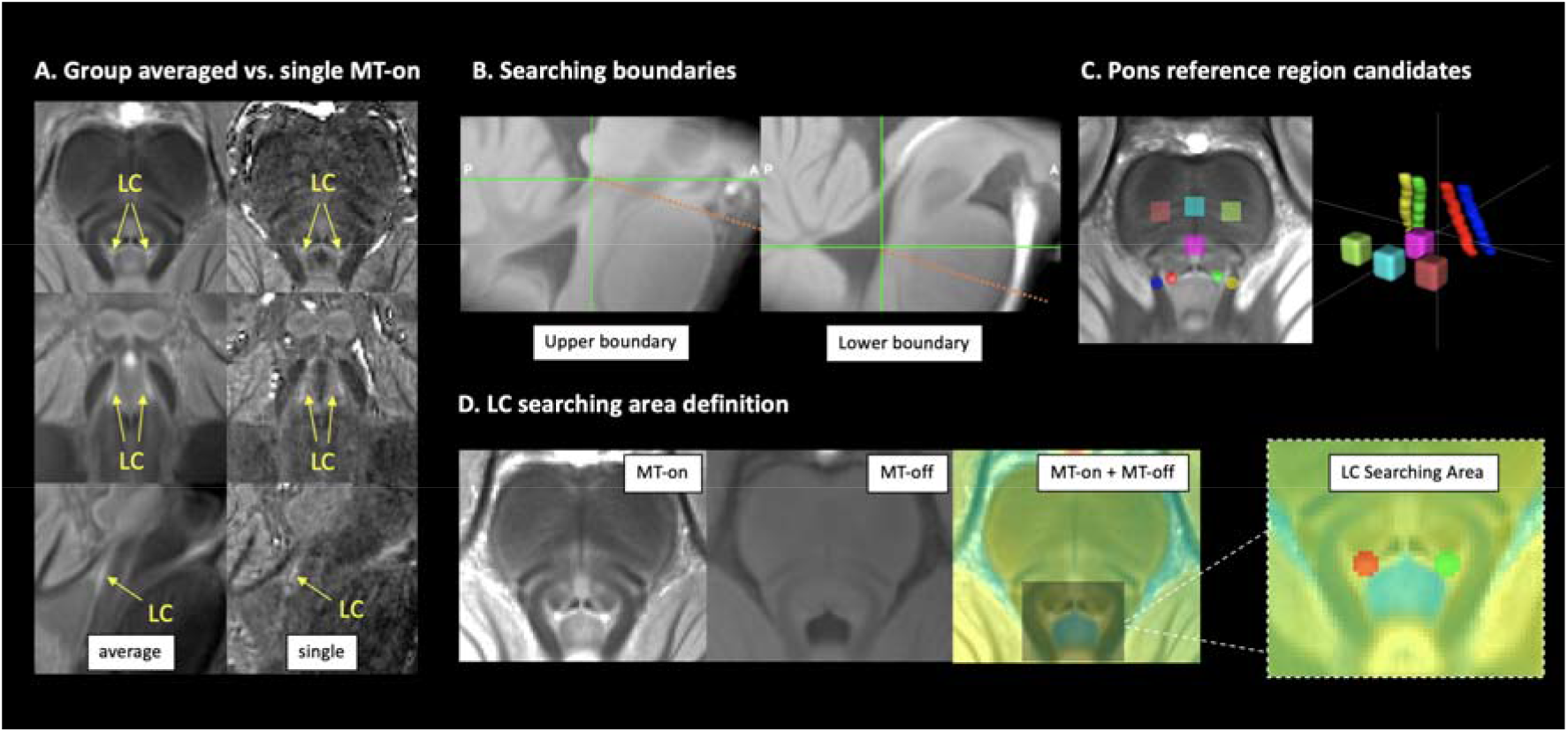
Anatomical definitions for LC segmentation. The LC was visible as the hyperintense regions on both group averaged and single MT-on images (A). To segment the LC using an unbiased and reliable semi-automated, threshold-based method, the upper and lower boundaries for searching the structure were defined at the level of the inferior colliculus and the recess th of the 4^th^ ventricle, respectively (B). Multiple brainstem areas were also defined for testing the most appropriate regions to be referenced to (C). Left and right LC searching areas were manually drawn on the axial view of the MT group template by overlaying the coregistered MT-on and MT-off images to avoid contaminations from voxels in the ventricle (D).

### 2.7 Probabilistic Atlas

For each axial slice on the rostro-caudal extent, the location of the left and right LC were estimated by representative coordinates of voxels with peak contrasts on each hemi-pontine region (Keren et al., 2009). The coordinates were then extracted for all subjects to form the group level spatial distribution of the LC and projected in a 3-D space for visualisation. Group standard deviations (SD) per axial slice were also calculated for X and Y coordinates to quantify individual variations. After applying the threshold for LC segmentation, the numbers of left and right LC voxels on each axial plane were counted to estimate the volumetric variation. Individual variations of peak CNR were also examined slice-by-slice, providing an independent contrast estimation from the LC volume which is inconsistent across subjects for valid comparisons.

A probabilistic atlas was created by adding and averaging the individual LC binary mask segmented from previous thresholding step. This yielded a map covering the spatial extent of the structure at the population level and capturing the between-subject variability indexed by the probability value for each LC voxel. The mean, median, and maximum probabilities were calculated for each axial slice separately. Multiple versions of the probabilistic atlas were produced with variable thresholds ranging from less to more stringent spatial definitions of the LC.

This new 7T atlas of the human LC was next compared with previously published atlases of the LC developed from 3T MRI (Keren et al., 2009, Tona et al., 2017, Betts et al., 2017). The MNI atlas from Betts et al. (2017) was downloaded from the source provided by the research group. We selected the 1SD version for the atlas published by Keren et al. (2009). Tona et al. (2017) provide LC atlases from two scan sessions and segmentations. These have been averaged and combined into one image to facilitate comparisons. Moreover, previous studies used the 0.5 mm 2006 version of the MNI template for the registration target. This standard image has minor differences in the lower part of the brain comparing to the ICBM152 2009b version which is the registration destination we used. To account for this, the two versions of the MNI template were coregistered with both linear and non-linear steps following the same registration pipeline in the previous step. The estimated transformation matrix was applied to the 3T LC atlases to correct the subtle difference due to the template selection.

### 2.8 LC Metrics and Age Effects

The spatial distribution of the LC signals and the individual variability of LC locations were examined slice-by-slice along the rostrocaudal axis. On each axial slice per hemi-pons, the segmented LC voxel with the peak contrast was used to represent the LC location. The coordinate and the CNR were extracted for each peak LC voxel and then 3D reconstructed to assess the spatial variability of the LC. After applying the threshold for LC segmentation, the number of LC voxels was also calculated and plotted slice-wise for all subjects.

To quantify the LC contrast, a CNR map was firstly computed voxel-by-voxel on the MT-on image for each subject using the signal difference between a given voxel (V) and the mean intensity in the reference region (Mean_REF_) divided by the standard deviation (SD_REF_) of the reference signals 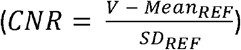. We compared the contrast against the SD instead of the widely used mean for calculating the CNR because the noise is more appropriately estimated by SD, as described in both signal detection theory and medical imaging analysis (Welvaert and Rosseel, 2013, Hendrick and Bradley, 1999). Peak LC CNRs were analysed slice-by-slice along the rostro-caudal extent. Mean LC CNRs were extracted by applying the probabilistic atlas with two variable thresholds to account for different levels of sensitivity-specificity trade-offs.

Previous work has reported age-related differences in LC contrast (Betts et al., 2017, Liu et al., 2019, Clewett et al., 2016) and suggested that the variation of the contrast during ageing was consistent with the concentration of neuromelanin in the LC (Zecca et al., 2004). Some studies also reported that age-related effects can confound the reference region selection by introducing biased baseline measures (Keren et al., 2009, Clewett et al., 2016). We thus investigated the effect of age with correlative models in Matlab (R2018b) to assess: 1) Pearson’s correlations between age and mean signal intensities within candidate reference regions to investigate whether the estimate of noise was confounded by age; 2) Pearson’s correlations between age and atlas-extracted mean LC MTR and CNR to test for age-related changes in MT signal and LC contrast; 3) canonical correlation analysis (CCA) (Sui et al., 2012) to test for age effect interacting with LC spatial variability on the rostro-caudal axis. We chose the CCA method as it provides latent variables with linear combinations for testing the relationship between age and individual spatial variability in the LC.

## 3. Results

### 3.1 Participants and MT sequence

There were no differences in the demographic characteristics of the two datasets (Table 1) which are aggregated for further analyses. Visible contrast and anatomical boundaries of the LC could be observed in all individual MT-on images, which were further enhanced on the group averaged MT-on template (Figure 2A).

**Table 1.** Demographic Summary of the Studies

|  | Total | Dataset 1 | Dataset 2 | Comparison* |
| --- | --- | --- | --- | --- |
| **Number** | 53 | 29 | 24 |  |
| **Age** | | | | $t(2, 51) = 0.93$<br>$p = 0.36$ |
| Mean (SD) | 66 (7.1) | 67 (8.2) | 66 (5.5) |  |
| Range (min/max) | 52/84 | 52/84 | 53/78 |  |
| **Gender** | | | | $\chi^2 = 0.005$<br>$p = 0.94$ |
| Male | 29 | 16 | 13 |  |
| Female | 24 | 13 | 11 |  |
| **MoCA** | | | | $t(2, 51) = -1.03$<br>$p = 0.31$ |
| Mean (SD) | 28.4 (1.4) | 28.2 (1.4) | 28.6 (1.4) |  |
| Range (min/max) | 24/30 | 24/30 | 25/30 |  |

\*demographic difference between studies were tested using two-sample t-test for Age and MoCA scores, and chi square test for Gender difference. MoCA: Montreal Cognitive Assessment

### 3.2 Registration Accuracy

As shown in Figure 1, individual MT images were warped to the standard MNI template with varied linear and non-linear parameters from the T1-driven co-registration pipeline. As critical step in the current study, the registration accuracy determined the consistency of LC segmentation and the interpretability of the results. To evaluate the registration accuracy most relevant to the LC analysis, pairwise DSCs were calculated between individual pons and the standard pons as an index for boundary similarity measures. The DSCs were very high across all subjects (mean ± SD: 0.96 ± 0.01, range: 0.9 – 0.97), suggesting that the co-registration pipeline was successful.

### 3.3 Reference region selection

Mean signal intensity, SNR, and MTR for LC searching area and PT reference candidate regions are plotted in Figure 3. Higher signals were consistently detected in the right SCP and PT areas relative to the left counterparts (Right SCP > Left SCP: t (52) = 7.12, *p* = 3.32 x 10, Right PT > Left PT: t (52) = 5.27, *p* = 2.68 x 10^−6^), whereas the mean signal intensities within the LC searching area did not differ between hemi-pontine regions (Right LC vs. Left LC: t (52) = 0.99, *p* = 0.33). This right-lateralised signal distribution indicated that the SCP and PT areas were not appropriate as reference regions, so the ipsilateral referencing strategy was not adopted as indicated in our a priori criteria. When considering the central and dorsal PT areas, the central PT showed high SNR (t (52) = 3.17, *p* = 0.0026) and MTR (t (52) = 14.77, p = 3.03 x 10^−20^), indicating that this ROI was an ideal candidate reference ROI for LC imaging analysis using MT-weighted sequences. Of note, the age-related signal reduction in the central PT area reported in previous studies using T1-w sequence was not observed in the current study (r = 0.05, *p* = 0.72).

**Figure 3.**
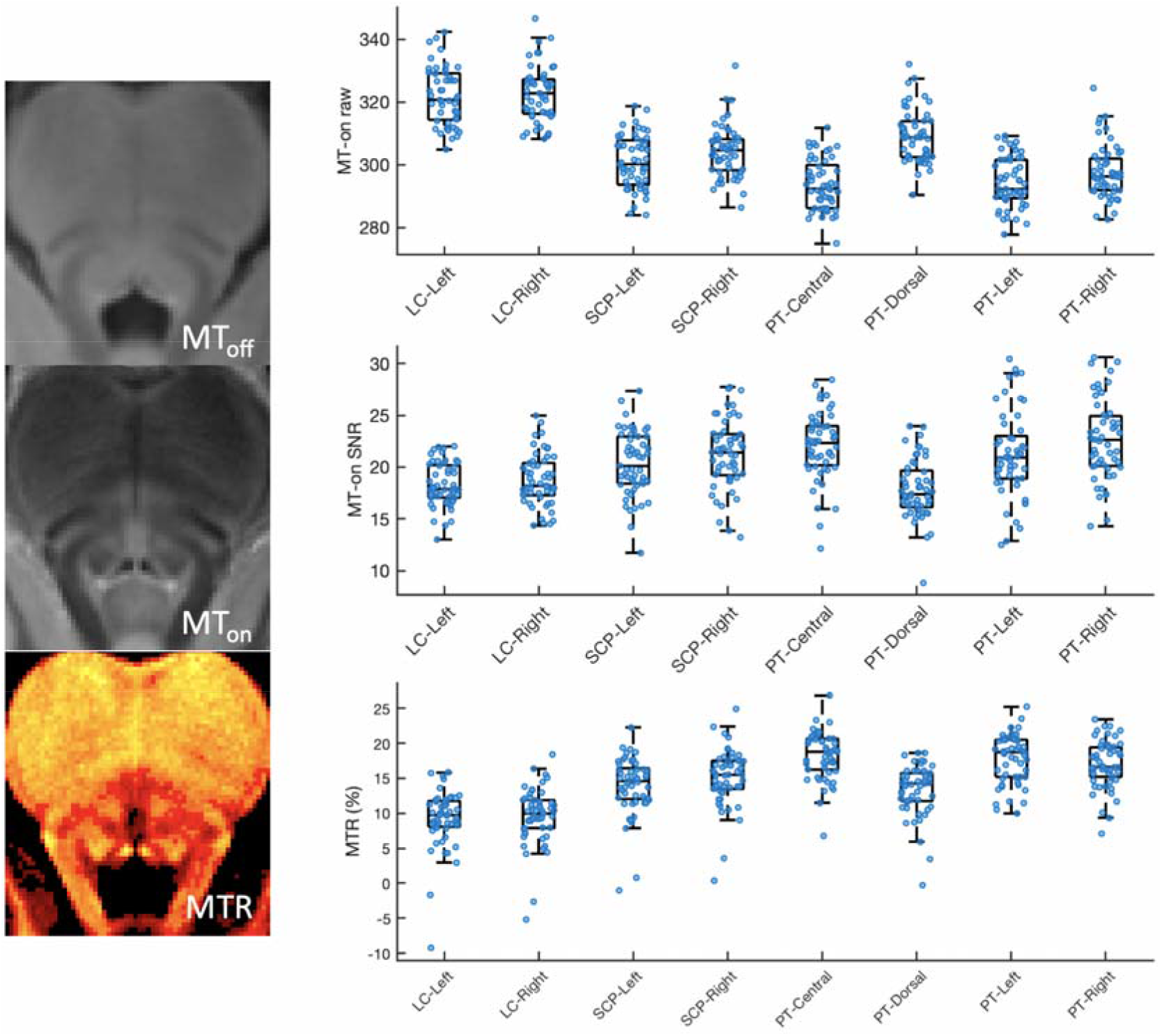
Signal evaluation for LC searching areas and candidate reference regions. The raw signals, the signal-to-noise ratios (SNR) and the magnetisation transfer ratios (MTR) for each region of interest were plotted. Scatter and box plots were both presented.

### 3.4 Spatial variability of the LC

Following the threshold-based, semi-automated segmentation pipeline, the LC was segmented in each normalised MT-on image for each subject by applying a threshold calculated from the central PT reference region. Slice-wise MNI coordinates of peak LC voxels from both hemi-pons were extracted and 3-D reconstructed as shown in Figure 4A. Along the rostro-caudal axis of the LC, lower degree of spatial agreement across subjects was observed in the caudal section. This is mimicked in the slice-wise SD plot showing the highest spatial variability with X and Y coordinates in the caudal part of the LC (Figure 4B).

**Figure 4.**
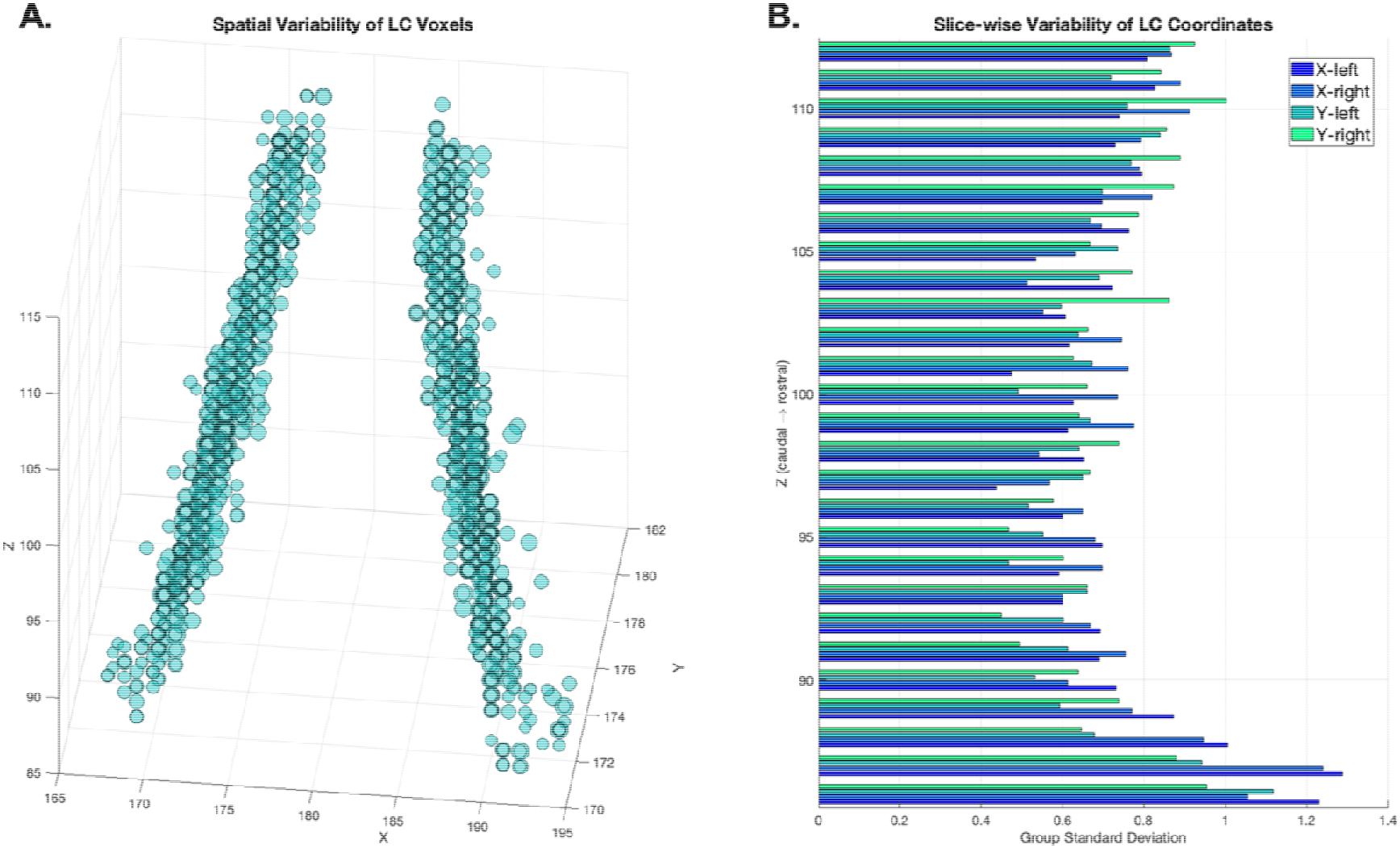
Spatial distribution and variability of LC voxels. (A) LC coordinates from all subjects were projected as circles in a 3D space showing the group variability of the spatial location of the LC. The coordinates were determined bilaterally on axial planes as the LC voxel with the highest contrast after threshold-based segmentation (> 5 SD). For each coordinate, the size of the circle corresponded to its degree of the contrast to noise ratio. (B) Group level spatial variability was demonstrated as the standard deviation of the X and Y coordinate along the rostrocaudal axis.

Slice-wise volumetric measures of the LC at 7T further revealed a distinctive spatial pattern for LC subregions. At the group level along the rostral-caudal axis of the LC, the number of LC voxels segmented using the thresholding method are shown in Figure 5.

**Figure 5.**
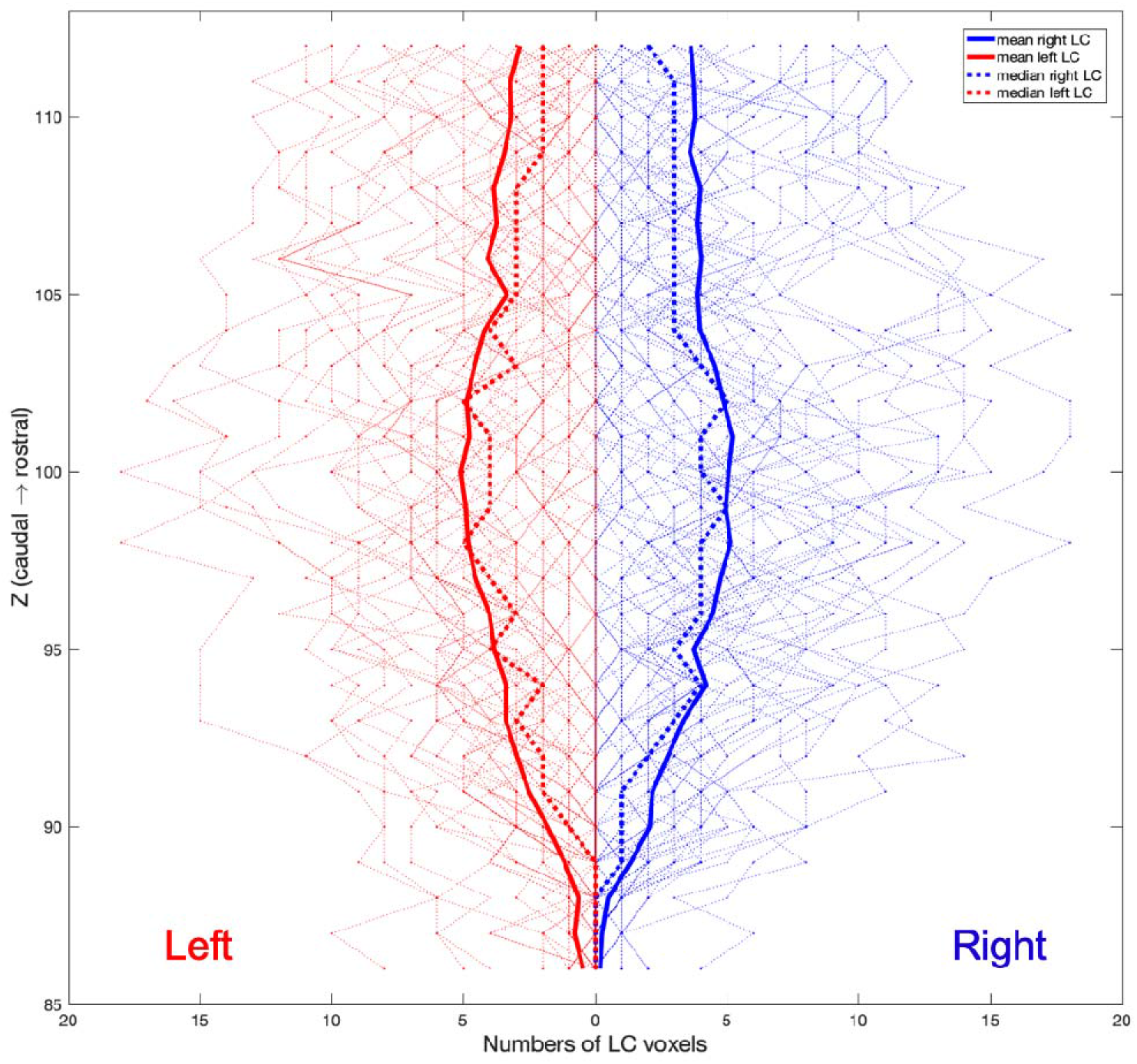
Individual and group level slice-wise voxel counts of left (red) and right (blue) LC. After the threshold-based segmentation (> 5 SD), LC voxels were counted for each subject and were plotted as the faded lines for each axial slice in the background. The group level mean and median were presented as the solid and dotted lines.

### 3.5 LC probabilistic atlas

A population-based probabilistic atlas of the LC was obtained by averaging individual LC binary masks. The maximum probability across all LC voxels was 86.79%. Slice-wise mean, median, and maximum probabilities are reported in Figure 6A. Two versions of the LC probabilistic atlas were created based on the slice-wise distribution of group-level probabilities. A liberal threshold (5%) was applied to obtain a template (705 voxels, 88.125 mm^3^) which is more sensitive to the LC spatial distribution while controlling for the potential noise seen in the caudal part of the LC. A stringent threshold (25%) was applied to the atlas to produce a template (284 voxels, 35.5 mm^3^) that is more specific to the core LC locations and which represent the highest mean probability across the whole rostro-caudal extent. The shape and spatial differences between the two types of the atlas are shown in Figure 6B.

**Figure 6.**
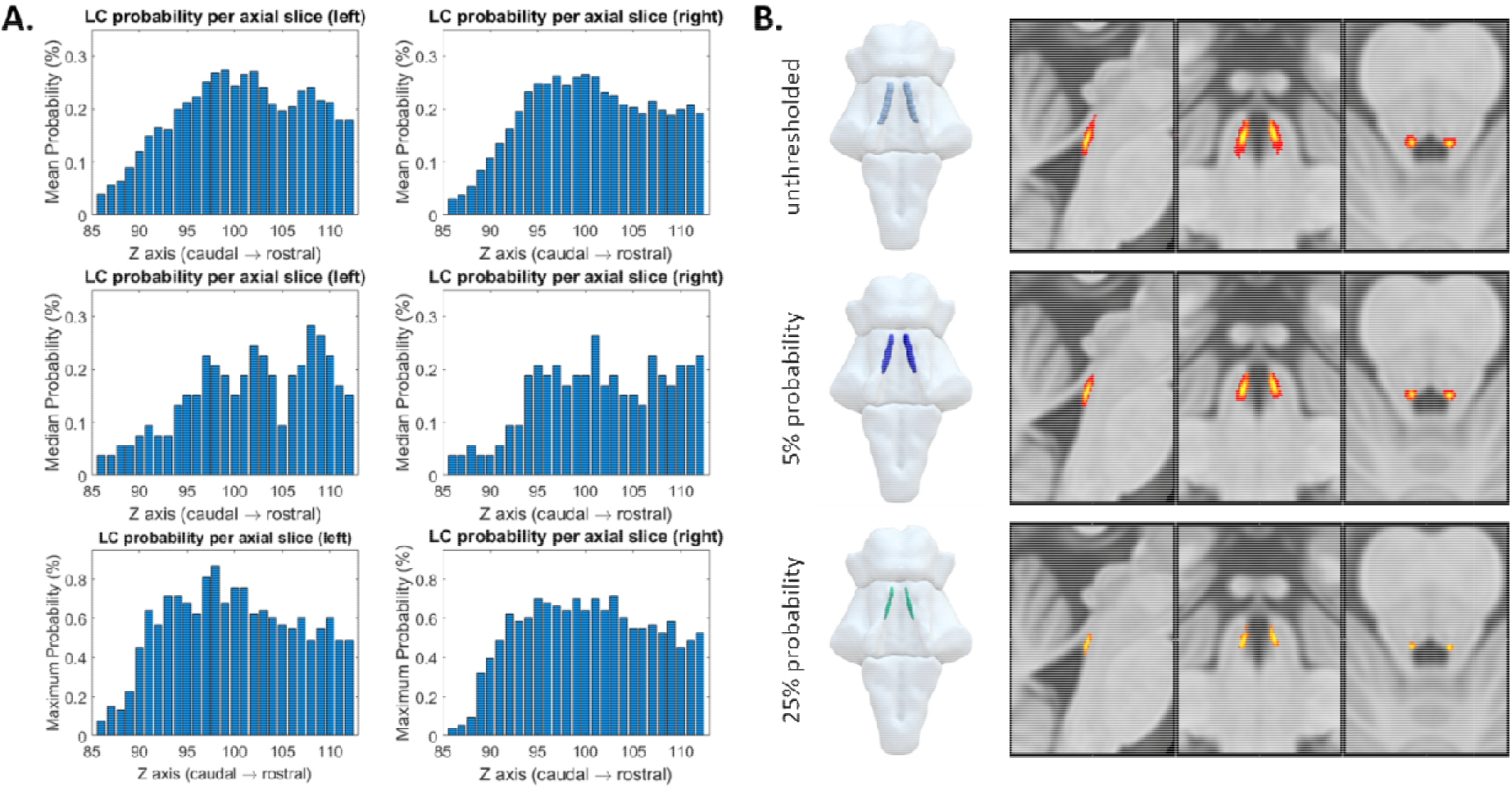
Probabilistic maps of the LC. (A) The mean (top row), median (middle row) and maximum (bottom row) probability per axial slice for left and right LC voxels. (B) The probabilistic LC maps thresholded at 0% (top), 5% (middle) and 25% (bottom) presented in 3D reconstructed brainstem and the multi-planar view overlaid on the standard ICBM152 0.5 mm T1-weighted image.

### 3.6 Comparison with published LC atlases

The 5% probability LC template was then compared to the published 3T atlases of the LC. The differences in imaging protocols and the spatial features are summarised in Table 2. In contrast to the 3T atlases, the 7T template is less extended at the caudal end and has more specific spatial boundaries for defining the LC (Figure 7).

**Figure 7.**
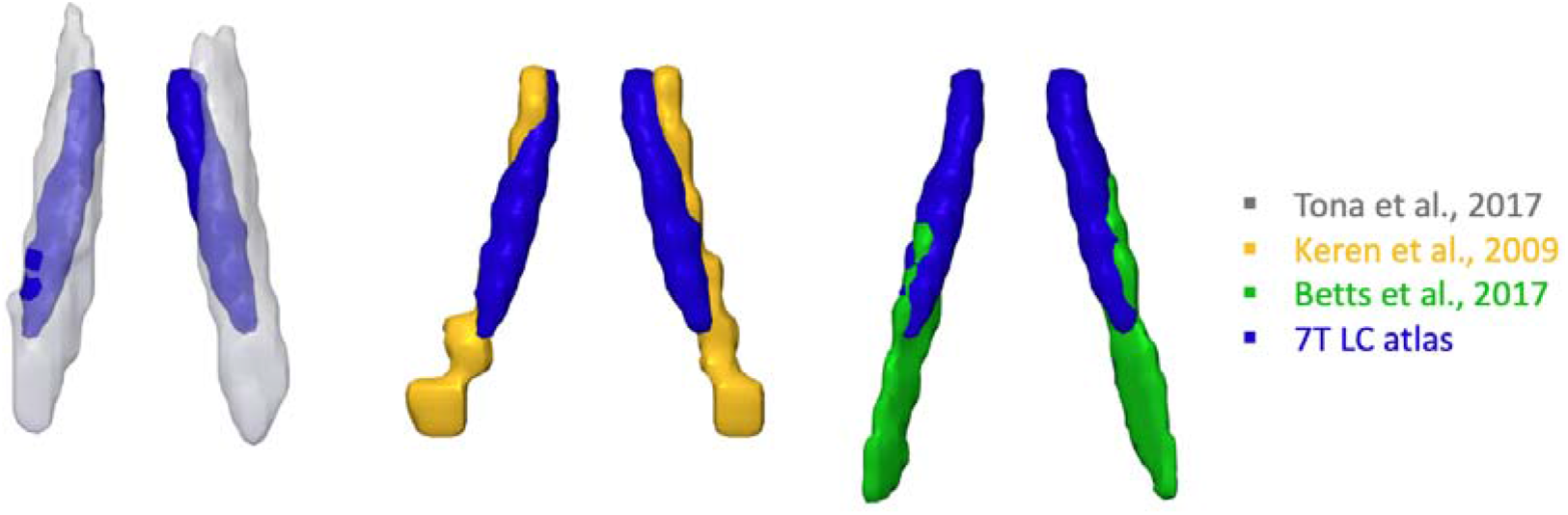
3-D renderings of 3T LC atlases from literature and the 7T LC atlas generated in this study (threshold 5%).

**Table 2.** Imaging protocols and atlas comparisons of 3T and 7T LC studies

| study | Method | N | Age (years) | Sequence | in-plane resolution (mm) | Thickness (mm) | Voxel volume (μl) | Coverage | Segmentation method | Variability map | Volume ($\text{mm}^3$) | Extent (Z in MNI) |
| --- | --- | --- | --- | --- | --- | --- | --- | --- | --- | --- | --- | --- |
| Keren et al. | 3T | 44 | 19 - 79 | T1-TSE | 0.4 x 0.4 | 3 | 0.48 | pons | manual peak | standard deviation | 98.12 | 113 - 77 |
| ona et al. | 3T | 17 | 19 - 24 | T1-TSE | 0.35 x 0.35 | 1.5 + 10% gap | 0.2 | pons | manual | probabilistic | 298.2 | 118 - 77 |
| etts et al. | 3T | 25<br>57 | 22 - 30<br>61 - 80 | T1-FLASH | 0.75 x 0.75 | 0.75 | 0.42 | whole brain | manual | none | 101.62 | 103 - 72 |
| 7T LC atlas | 7T | 53 | 52 - 84 | MT-TFL | 0.4 x 0.4 | 0.5 | 0.08 | pons + midbrain | semi-automated | probabilistic | 88.13 (5%)<br>35.5 (25%) | 112 - 86 |

### 3.7 LC CNR distributions and age effects

Slice-wise CNRs were extracted for LC voxels with peak contrast or by implementing the atlas-based method for LC voxels within the probabilistic atlas to account for between-subject volumetric differences. On each axial slice, peak LC CNR and mean CNR calculated within the 5% and 25% probability atlases were plotted in Figure 8. Consistently with the LC volume measures (Figure 5), group mean peak CNR was increased gradually from the rostral end and reached to the maximum in the central section for both the left and right LC, then decreased in the caudal LC. However, this rostro-caudal gradient of LC CNR was less evident using atlas-based method for CNR extractions.

**Figure 8.**
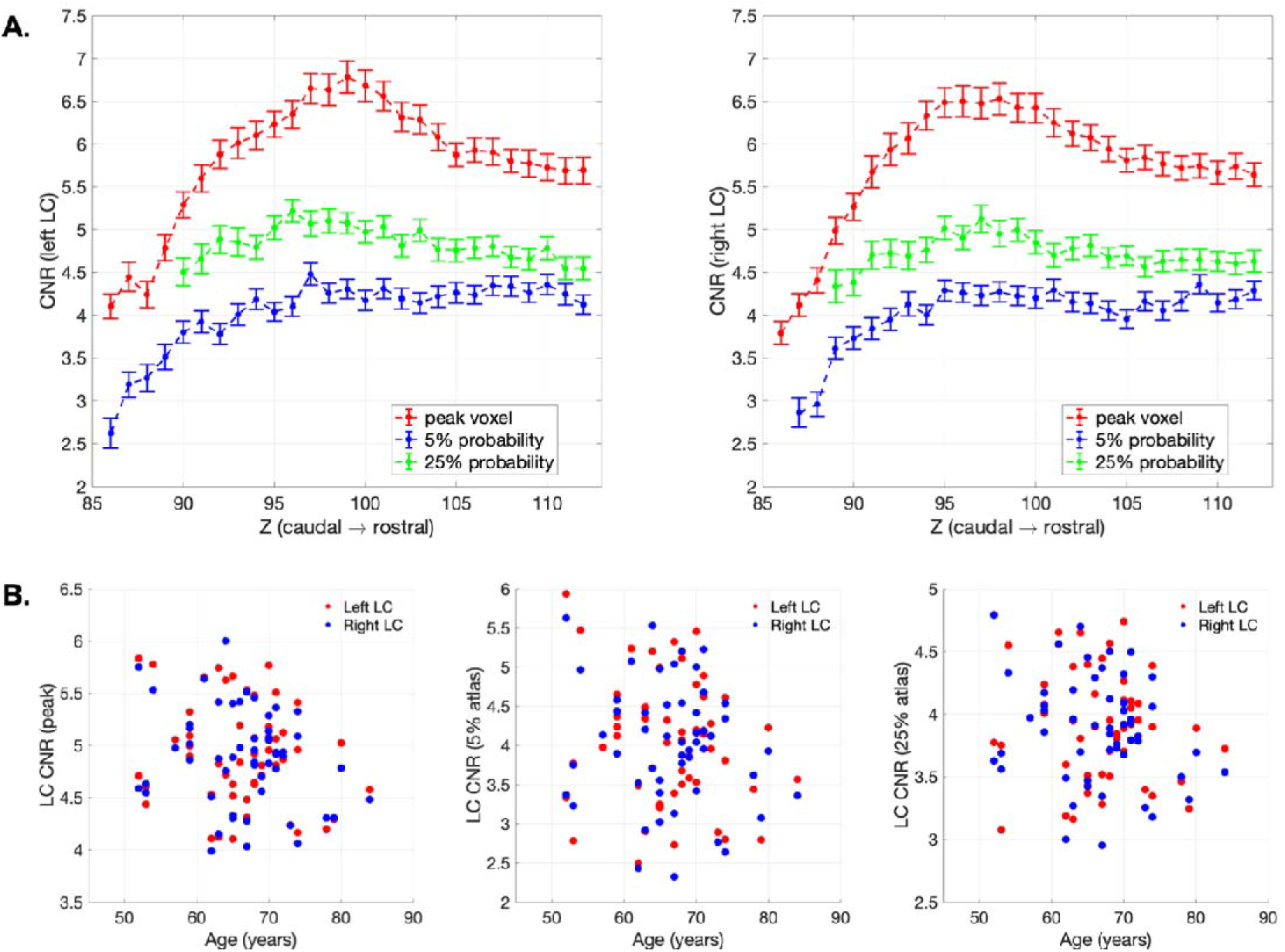
LC CNR and the correlation with age. (A) peak and atlas extraction of the CNR for left and right LC per axial slice. Standard error bars were presented. (B) scatter plots demonstrated the relationships between age and peak- and atlas-extracted CNRs.

There was no significant age effect on either overall or slice-wise LC CNR measures (Figure 8) examined using Pearson’s correlation and CCA analyses (p > 0.05).

## 4. Discussion

In this study, the details of the LC spatial features have been revealed using a sensitive high-resolution MT-weighted sequence at ultra-high field. This characterization of the LC anatomy and the contrast sensitivity across the structural distribution of the LC support an *in vivo* probabilistic atlas. This 7T MRI atlas has promising applications in future studies investigating the structural or functional changes of the LC in neurodegenerative disorders. The anatomical localisation of the LC in the current study benefits from several advantages.

First, a high-resolution definition of the LC was achieved using the MT-TFL sequence and the 7T MRI. The SNR-resolution trade-off is critical due to the small size and the challenging anatomical location of the LC. Previous 3T MRI studies have needed to sacrifice resolution to prioritise sufficient SNR for detecting the LC. The 7T MT-weighted sequence adopted here resolved this conflict and provided comparable SNR to the 3T T1-weighted sequence while enabling a 3-D high-resolution imaging that accounts for partial volume effects (Priovoulos et al., 2018). Robust LC contrast was directly elicited by the MT saturation pulses, possibly due to the increased saturation of the tissue surrounding the LC, as shown by a series of recent studies (Trujillo et al., 2017, Watanabe et al., 2019a, Watanabe et al., 2019b, Trujillo et al., 2019).

Second, the quality of our LC atlas was improved by tailored registration and segmentation methods. An MNI template was used for spatial normalisation and to minimise inconsistencies in segmentation and normalisation. Highly accurate registration specific to the pons (96% on average) was achieved following the T1-driven diffeomorphic co-registration pipeline. A semi-automated segmentation protocol with excellent reliability was then applied to extract individual LC voxels based on a predefined threshold (Langley et al., 2017, Dahl et al., 2019). This pipeline was specifically designed to minimise the inevitable biases of manual segmentation and to increase reproducibility of results via the use of the same dataset and scripts. The selection of the most appropriate reference region for segmentation and contrast calculation resulted from a systematic comparison of the signal metrices across 6 candidate regions. The central pontine area was chosen because of its high SNR and MTR. We also detected a lateralisation of the signal intensity for pontine areas but not for the LC. This suggests that left and right pontine regions need to be used with caution as reference regions as they can introduce biases during the comparison between the left and right LC contrast.

Third, amongst the registration and segmentation steps, there were only two elements that required user interventions, i.e. the definition of the LC searching volume and the determination of the threshold. The circular searching area was defined on the axial plane with 2.5 mm diameter similar to the width of the LC reported in *ex vivo* studies (German et al., 1988, Fernandes et al., 2012). A more stringent 5 SD instead of the 4 SD (Chen et al., 2014, Dahl et al., 2019) threshold was used for LC segmentation to account for the enhanced contrast at *7T*. Significant displacements were found along the rostro-caudal axis of the LC across previously published atlases and this is likely due to disagreements regarding the superior and inferior boundaries of the LC (Table 2 and Figure 7). The rostral part of the LC in this study was defined (in the MNI space) at the level of the inferior colliculus, which is approximately located at the same level of the superior boundary of the pons (Figure 2). This is consistent with the anatomical definition of the LC in the majority of previous imaging and post-mortem studies (Keren et al., 2009, Tona et al., 2017, German et al., 1988, Fernandes et al., 2012). In contrast, the caudal part of the LC has been found to terminate at various locations across different neuroimaging studies. This variance probably reflects ‘real’ individual variations of the LC length but may also have resulted from partial volume effects deriving from the use of thick slices. Our data show a possible LC extent of ~15.5 mm along the brainstem long axis which accords with the average ex *vivo* measurement of 14.5 mm (Fernandes et al., 2012).

Fourth, the characteristic spatial features were identified after an automated LC segmentation with striking similarities relative to *ex vivo* findings and biophysical measures. A rostro-caudal gradient was consistently observed for LC volumes (Figure 5), contrast signal (Figure 8), spatial variance (Figure 4), and the percentages of overlap (Figure 6) across all 53 subjects. The volumetric and contrast reduction from the rostral to caudal parts of the LC mimicked the spatial pattern of LC cell number and density reported in previous *post mortem* research (German et al., 1988, Fernandes et al., 2012). Similar spatial distribution was also observed at the level of macromolecular content, as indexed by quantitative MT measurement on the pool size ratio (Trujillo et al., 2019). Of note, the central segment of the LC showed the lowest between-subject spatial variability in this imaging study and in a previous stereological *post mortem* work (Fernandes et al., 2012). The estimated volumes of the whole and core LC in another *ex vivo* study (whole: 71 mm^3^, core: 36.7 mm^3^) were close to the current 7T LC atlas thresholded at different probabilistic levels (5%: 88.13 mm^3^, 25%: 35.5 mm^3^) (Tona et al., 2017).

We found a discrepancy between the peak CNR and the atlas-based CNR in the central segment of the LC relative to its rostral and caudal parts (Figure 8). In general, the overall contrast difference across variants of the CNR reflected the level of representativeness associated with the signal density when applying different sizes of masks for extracting the mean contrast. We propose that the regional contrast difference in the central LC with peak or atlas extractions can be explained by the distinctive spatial distribution of the LC neurons within the central area. Considering that the LC cells are not uniformly distributed on both the axial plane and rostro-caudal axis, the LC CNR extracted from single voxels might be inherently susceptible to the interaction of cell density and cell number as it only measures the contrast originated from the core LC. Fernandes et al. (2012) reported that the LC cells were most closely organised in the central LC and more dispersed in the rostral and caudal ends. In addition, German et al. (1988) reported a non-linear reduction of the LC cell count from the rostral to caudal part of the LC where the numbers of rostral and central LC cells were indifferentiable. The increased CNR peak in the central LC observed here might therefore reflect a greater amount of LC neurons populating this area, whereas the CNR extracted via the 7T atlas within a much bigger area may be less sensitive to the local cell density which may be more directly associated with the absolute cell number.

Finally, we examined the effect of age on the LC contrast. The impact of ageing on LC signal has been previously been assessed (i) by showing the group difference between young and older adults (Clewett et al., 2016, Betts et al., 2017, Dahl et al., 2019), or (ii) by testing for a linear and quadratic function of age across the life-span (Shibata et al., 2006, Liu et al., 2019). The results are consistent with the accumulation of neuromelanin with age (Zecca et al., 2004). In this study, we did not find a significant age-related difference in the LC CNR, perhaps because of the narrower age range (52-84 years). We used a relatively narrow age-range in the view of the use of the atlas for studies of disorders associated with later life. However, it is also worth noting that, although a non-linear relationship between age and LC contrast may occur (Zucca et al., 2014, Sulzer et al., 2018), ageing in itself is a complex process which may not necessarily be a reliable predictor of individual variability in the LC integrity.

Several limitations on the interpretation and application of the atlas need to be addressed here. First, the specific source of the contrast in the LC using either T1-weighted or MT-weighted sequence is still unclear, hindering the establishment of a meaningful biophysical model for the contrast. Post mortem and animal work has confirmed that the presence of noradrenergic LC neurons is prerequisite for the detection of enhanced contrast in the LC (Keren et al., 2015, Watanabe et al., 2019a). Previous studies have linked the contrast to the T1 reduction and the MT effect contributed by the intraneuronal neuromelanin macromolecules (Chen et al., 2014, Trujillo et al., 2017, Priovoulos et al., 2018, Sulzer et al., 2018). However, contrast in the LC may also partially originated from the enriched water content (Watanabe et al., 2019a, Watanabe et al., 2019b). Further work is required to quantify and dissociate these mechanisms as contributors to human LC contrast. Second, the clinical application of the atlas-based estimates is constrained by variability in the pathological changes in the LC, some of which may exert a stronger influence on the MT-weighted contrast e.g. neuronal death, neuronal inclusions and changes in neuronal volume in early stages (Theofilas et al., 2017). Moreover, the role of pathology-related events such as neuroinflammation in the formation of MR properties of the LC is also unclear (Lambert et al., 2013, Watanabe et al., 2019a) but directly relevant to several neurodegenerative disorders (Passamonti et al., 2018b). Third, MR artefacts may occur with ultra-high resolution due to the flow effect from blood vessels and head movements. Further optimisation of the sequence for imaging the LC might include flow compensation (Haacke and Lenz, 1987) and explicit motion correction with prospective or retrospective techniques (Lusebrink et al., 2017, Schulz et al., 2012, Stucht et al., 2015).

The thresholded atlases are freely available at https://www.nitrc.org/projects/lc_7t_prob/. The 53-subject MT and MP2RAGE data used in this study are also fully available for noncommercial academic purposes upon request.

## 5. Conclusion

This study provides a comprehensive 7T probabilistic atlas of the human LC at the finest resolution to date, in order to facilitate the *in vivo* localisation of this important brainstem nucleus. A robust rostro-caudal gradient of the LC spatial features was observed using sensitive MT-weighted images from 53 healthy elderly individuals in combination with accurate registration and reproducible segmentation. The LC metrics extracted using the methods described in this study are highly consistent with previously published *ex vivo* findings. The LC contrast estimation, unbiased by volumetric differences, will be a useful tool in clinical studies for detecting *in vivo* pathological changes in the LC.

## Acknowledgements

This work is supported by the UK Medical Research Council (MR/P01271X/1), the Wellcome Trust (103838), the British Academy (PF160048), the China Scholarship Council (CSC), a Neil Hamilton Fairley Fellowship from the National Health and Australian Medical Research Council (#1091310), a Sir Henry Dale Fellowship from the Wellcome Trust and the Royal Society [098436/Z/12/B], a Lundbeckfonden International Postdoc Fellowship (R232-2016- 2333/R265-2017-3722), a Cambridge Trust Vice-Chancellor’s Award and Fitzwilliam College Scholarship, NIHR Cambridge Biomedical Research Centre and the Cambridge Centre for Parkinson-plus. The views expressed are those of the author(s) and not necessarily those of the NHS, the NIHR or the Department of Health and Social Care.

